# MAGNET: an all-in-one foundation model for cross-modal and cross-dimensional microscopic image restoration

**DOI:** 10.64898/2025.12.23.696141

**Authors:** Yifan Ma, Tianfeng Zhou, Lanxin Zhu, Chengqiang Yi, Yunshi Zhou, Binbing Liu, Peng Fei

**Affiliations:** School of Optical and Electronic Information-Wuhan National Laboratory for Optoelectronics, Huazhong University of Science and Technology, Wuhan, China, 430074; Department of Ophthalmology, Tongji Hospital, Tongji Medical College, Huazhong University of Science and Technology, Wuhan, 430030, China; Advanced Biomedical Imaging Facility Huazhong University of Science and Technology, Wuhan, Hubei 430074, China

## Abstract

The application of emerging Foundation models in current microscopy has remained limited due to task-specific model designs, strong modality dependence and insufficient robustness under complex degradations. Here, we introduce MAGNET (Microscopic All-in-one General fouNdation model for imagE resToration), the first end-to-end All-in-One foundation model designed for universal microscopic image restoration. MAGNET integrates multi-task, cross-modal and cross-dimensional (2D + 3D) image restoration capabilities within a single unified architecture, and it is specifically designed to handle composite degradations in practical imaging systems. The framework comprises three components: (i) a task-aware, prompt-guided feature enhancement module for adaptive learning across diverse degradations; (ii) an LIIF-based reconstruction module enabling continuous, resolution-adaptive restoration; and (iii) a dimension-compatible triple-plane projection module supporting both 2D and 3D data. Trained on large-scale datasets spanning 5 major microscopy modalities (structured illumination, confocal, light-sheet, wide-field and two-photon) and 8 representative restoration tasks, MAGNET achieves state-of-the-art performance across super-resolution, denoising, deblurring, background suppression, isotropic reconstruction, aberration correction, de-scattering and virtual staining. Beyond supervised training, MAGNET supports direct inference on unseen systems, efficient few-shot fine-tuning and self-supervised test-time optimization, enabling robust performance across diverse data and imaging conditions. MAGNET serves as a unified, generalizable, and scalable computational framework for microscopy, enabling more efficient and reliable downstream analyses in cell biology, neuroscience, and pathology.

## Introduction

Fluorescence microscopy^1^ is an indispensable tool for modern biological research, providing molecularly specific access to cellular architecture, protein localization and dynamic physiological processes. Yet image quality is intrinsically degraded by a combination of physical and experimental factors^2^: the finite numerical aperture limits spatial resolution, the short exposure time required for high-speed imaging reduces the signal-to-noise ratio (SNR), and the scattering and refractive-index– induced aberrations arising in deep-tissue imaging impair signal quality. These factors obscure fine cellular structures and propagate errors into downstream analyses. Insufficient resolution can mask nanoscale features essential for studying organelle dynamics; noise and background fluorescence undermine the accuracy of quantitative measurements in live-cell imaging; and inconsistent fluorescence patterns complicate cross-sample normalization in pathology and high-content screening. Effective computational restoration is therefore indispensable for recovering high-fidelity biological information, supporting robust quantitative analysis and enabling reliable interpretation across diverse biological applications.

Over the past decade, deep learning has transformed microscopic image processing, driving advances in image reconstruction^3, 4^, denoising^5^, super-resolution^6^, registration^7^, segmentation^8^, and related low-level vision tasks. Combined with a wide range of imaging modalities—including structured illumination microscopy^9^ (SIM), confocal^10^, two-photon^11^, and wide-field(WF) microscopy^12^—these methods have improved image quality and facilitated reliable downstream biological analysis. Denoising methods^5, 13^ improve SNR under low-light or high-speed conditions, super-resolution approaches^6, 14^ break the diffraction limit to reveal subcellular structures, and fluorescence-to-Haematoxylin-and-Eosin (H&E) style transformations^15^ enhance morphological interpretability and link fluorescence imaging to pathology. Together, these developments have moved deep learning–based microscopy image processing from a collection of isolated technical improvements to an increasingly well-established methodological framework.

Despite this progress, most existing approaches still follow a task-specific design paradigm: they combine data from one or a few modalities with a known degradation model to construct task-tailored network architectures and training strategies. For example, DFCAN^16^ incorporates a Fourier -domain module to enhance reconstruction of SIM images by exploiting their characteristic periodic patterns; and SeReNet^3^ introduces a dedicated deblurring–fusion module for multi-angle light-field microscopy. Such designs depend strongly on assumptions about the underlying data distribution and typically perform well only under constrained settings, such as a single task or a fixed acquisition protocol, where the inverse problem is well approximated^17^. When exposed to imaging modalities outside the training distribution, composite degradations or structural variations, performance often degrades markedly, with poor transferability, weak generalization and high deployment cost that limit scalability in realistic biological imaging scenarios.

In computer vision, foundation models^18^ have emerged as a promising route toward more general-purpose systems, inspired by large language models^19^ (LLMs) in natural language processing. By pre-training on large-scale, multi-task, and multi-modal datasets, they learn general-purpose visual representations with strong transferability and adaptability. As a representative example for high-level vision tasks, CLIP^20^ aligns images and text in a shared embedding space to enable cross-modal understanding, whereas FoundIR^21^, designed for low-level vision tasks, achieves robust and generalizable image restoration by integrating a million-scale real-world restoration dataset with a hybrid generalist–specialist architecture. Despite their impressive performance, current foundation models^22, 23^ still exhibit limited generalization without task-specific fine-tuning, particularly when applied to distinct visual distributions or modalities. To overcome these limitations, a new trend of “All-in-One”^24, 25^ foundation models has emerged, aiming to unify multi-task and multi-modal learning through shared architecture and feature representation. While highly successful in natural image domains, transferring these models to microscopy remains challenging due to the intrinsic heterogeneity of imaging systems (e.g., SIM, confocal, light-sheet), limited data availability, the complexity of physical degradations, and the dimensional incompatibility across multi-dimensional data (2D + 3D). These domain-specific variations introduce significant distribution shifts and leading to catastrophic forgetting^26^, inter-task interference^27^, and severe performance drops on out-of-distribution data. UniFMIR^28^ represents the first attempt to introduce a foundation-model paradigm into fluorescence image restoration and has demonstrated the feasibility of cross-task learning in microscopy, achieving promising performance across multiple restoration tasks. However, its design still follows a task-specific paradigm: multi-task capability is achieved through separate task heads and tails, limiting scalability when encountering new degradations, modalities or composite imaging conditions.

To address these challenges, we developed MAGNET (Microscopic All-in-one General fouNdation model for imagE resToration)—the first end-to-end All-in-One foundation model specifically designed for microscopic image restoration. MAGNET integrates multi-task (8+ tasks), cross-modal (5+ imaging modalities), and cross-dimensional (2D + 3D) restoration within a unified framework, while explicitly accounting for composite degradations that arise in real imaging systems. Its architecture combines three key components: (1) a task-aware feature enhancement module based on X-Restormer, which employs content-adaptive prompts to modulate degradation-specific features and alleviate conflicts across heterogeneous imaging conditions and morphologies; (2) an LIIF-based continuous scale reconstruction module, which provides continuous scale and resolution-adaptive restoration for diverse imaging configurations; (3) a dimension-compatible triple-plane projection module, which enables unified representation of 2D and 3D data by integrating complementary spatial priors. Beyond supervised training, MAGNET adopts a progressive deployment paradigm—including direct inference on unseen systems, few-shot fine-tuning and self-supervised test-time optimization. This flexible design allows a single model to adapt to varying data availability and experimental constraints. Extensive evaluations across more than 10 public and in-house datasets show that MAGNET achieves state-of-the-art performance over a wide range of microscopy restoration tasks and imaging conditions, establishing a unified, generalizable and scalable foundation model for quantitative microscopic imaging.

## Results

### Overall pipeline of MAGNET

Microscopy data inherently span multiple dimensions, including 2D, 3D, and time-lapse sequences, which encompass diverse imaging tasks and complex degradations (Fig. 1a). These degradations include optical blurring from the finite numerical aperture of the imaging system, stochastic noise introduced by the detector, and aberrations due to refractive index mismatches between the objective lens and the specimen.

**Fig. 1.**
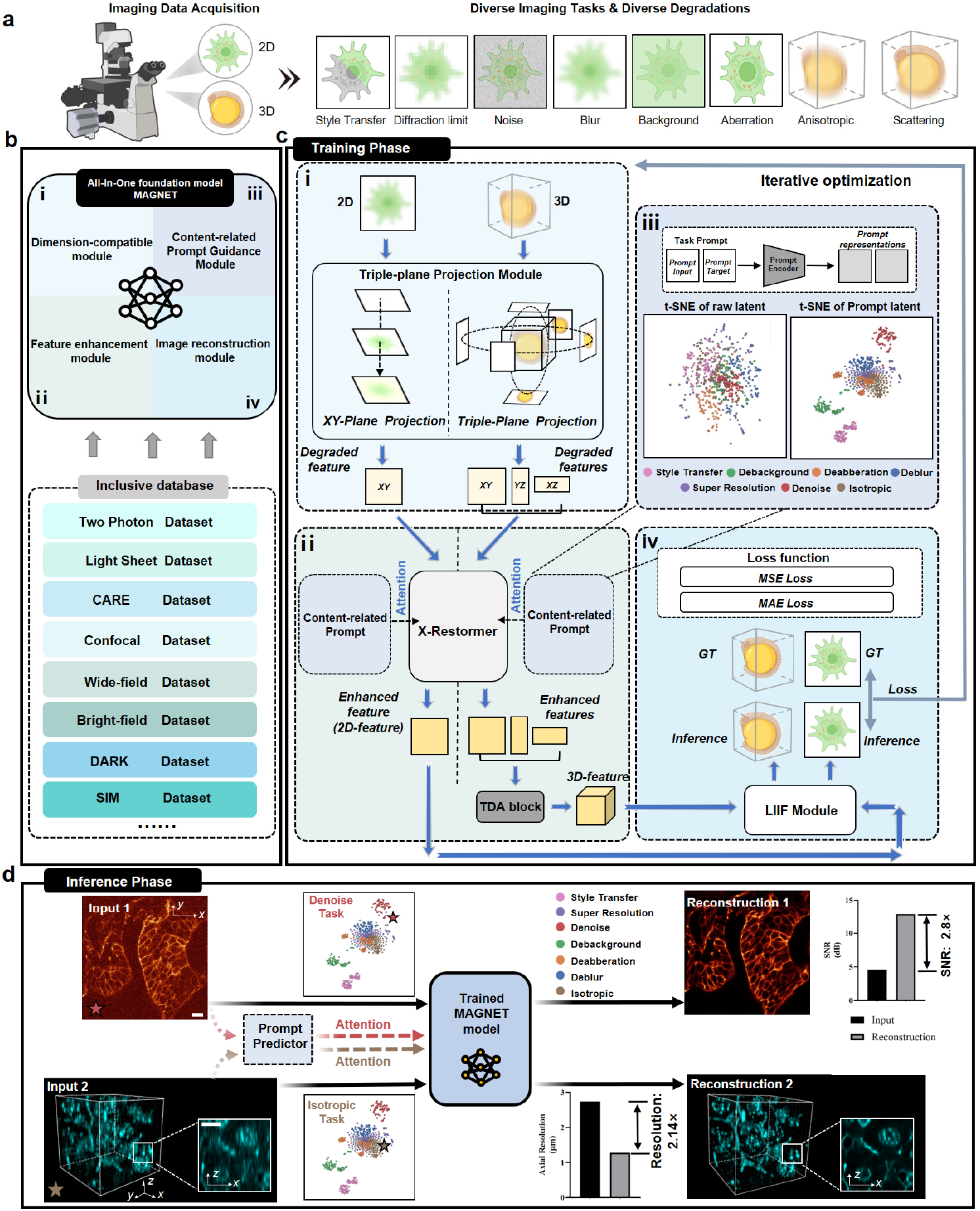
Overall pipeline and architecture of MAGNET for microscopic image restoration. (**a)** Microscopy data span multiple dimensions (2D, 3D, and time-lapse) and encompass diverse imaging tasks and degradations, including style-transfer, diffraction-limited resolution, noise, blur, background, aberration, anisotropy, and scattering. **(b)** Architecture of MAGNET integrating four modules: dimension-compatible module; feature enhancement module based on X-Restormer; content-related prompt guidance module; and a LIIF-based image reconstruction module. The model is trained on a large-scale heterogeneous microscopic image dataset covering major imaging modalities and biological scales. **(c)** Training workflow of MAGNET. **(i)** The dimension-compatible module (Triple-Plane Projection) unifies processing of both 2D and 3D data by projecting volumetric inputs along the x, y, and z directions to form complementary 2D representations. **(ii)** The feature enhancement module strengthens contextual representations across degradations and modalities through an X-Restormer backbone. **(iii)** The content-aware prompt guidance module dynamically modulates feature representations based on task-specific degradation patterns, improving task discriminability. The t-SNE display inside the figure shows the raw and prompt latent representations from MAGNET. The latent features extracted by the network was embedded in two-dimensions using t-SNE and clustered into seven clusters according to degradation labels. **(iv)** The LIIF-based image reconstruction module supports continuous scale reconstruction in both 2D and 3D. **(d)** Inference phase. The trained model employs a Task Prompt Predictor to automatically infer degradation-related prompts from input images and guide targeted restoration. On a noisy image of zebrafish embryos (Input 1), denoising prompts enable SNR enhancement (2.8× improvement), while on an anisotropic mouse liver volume, isotropic prompts enable axial resolution recovery (2.14× improvement, Input 2). Scale bars: 10 μm.

To restore these degradations with a unified model, we present MAGNET, an All-in-One foundation model for low-level image restoration tasks in microscopy (Fig. 1b). The framework introduces four complementary modules that jointly tackle the fundamental challenges of all-in-one learning: a dimension-compatible module for unified 2D/3D representation; a feature enhancement module based on X-Restormer; a content-related prompt guidance module for adaptive degradation-aware modulation; and a LIIF-based image reconstruction module enabling continuous and resolution-adaptive restoration across modalities. Together, these modules establish a unified architecture that balances shared representation learning with task-specific modulation—a key requirement for effective all-in-one restoration. To enable robust All-in-One learning and allow the model to capture diverse degradation characteristics in microscopy, we constructed a large-scale, heterogeneous microscopic image training dataset by consolidating data from diverse sources (Supplementary Table1). This dataset encompasses major imaging modalities—such as confocal, WF, SIM, light-sheet, and two-photon microscopy (including all aforementioned restoration tasks and degradation types)—and covers biological targets ranging from organelles (e.g., mitochondria, nuclei) to tissue fragments (e.g., Drosophila embryos, zebrafish). It spans structural scales from subcellular level to tissue levels and includes both 2D images and 3D volumetric data, representing a wide range of experimental conditions and restoration requirements (Fig. 1b).

Specifically, in the training phase (Fig. 1c), the model sequentially processes microscopy data through four core modules. First, the dimension-compatible module (Triple-Plane Projection) (Fig. 1ci, Methods) enables unified processing of both 2D and 3D data by integrating the priors inherent to each dimensionality. For 3D volumetric data, this module projects the volume along the x, y, and z axes to generate complementary 2D initial feature maps, facilitating the subsequent network module to efficiently process information from different perspectives. Second, the extracted initial features are refined by a feature enhancement module based on X-Restormer backbone, which strengthens contextual representations across diverse degradations and imaging modalities. (Fig. 1cii and Supplementary Fig .1) For 3D data, a dedicated reconstruction module then fuses the multi-view 2D projection features back into the original 3D volumetric representation, ensuring spatial consistency and completeness. Third, within this backbone, a content-aware prompt-guidance mechanism (Fig. 1ciii, Methods) adaptively modulates feature representations according to task-specific degradation patterns. By explicitly separating task-relevant information, this module mitigates the interference commonly observed in conventional multi-task models. Supporting t-SNE analyses show that prompt guidance leads to well-separated and more discriminative task clusters (Supplementary Fig. 2), confirming the mechanism’s effectiveness in mitigating inter-task confusion. Finally, the model incorporates a LIIF-based image reconstruction module (Fig. 1civ, Methods), whose continuous-scale representation fitting enables unified modeling and reconstruction of microscopy images across varying pixel resolutions and imaging configurations, thereby enhancing adaptability in multi-task microscopy scenarios. The detailed schematic of the network architecture is provided in the Supplementary Fig .3.

**Fig. 2.**
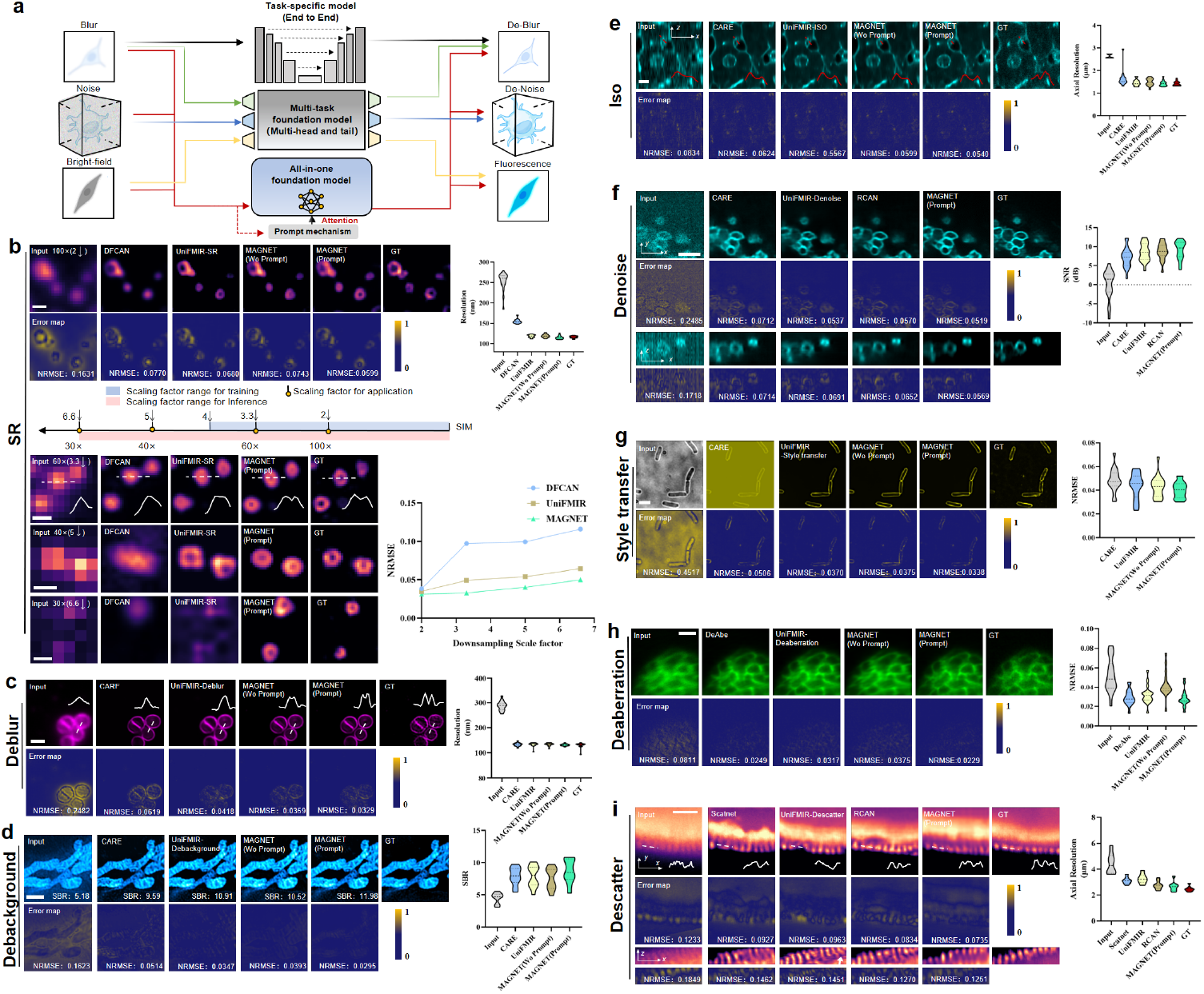
Multi-task All-in-One restoration of cross-dimensional fluorescence microscopy data. **(a)** Training schematic of existing task-specific and multi-head/tail foundation models and the MAGNET framework. Task-specific models focus on single degradations, multi-head/tail foundation models rely on separate branches for multi-task training, and the MAGNET employs a shared architecture with prompt-guided modulation to perform multi-task restoration. **(b)** Super-resolution (SR): Examples of the SR performance of DFCAN, UniFMIR-SR, and MAGNET (with/without prompt) on CCPs samples from BioSR dataset; The NRMSE*↓* (lower is better) values are shown on the error map images under the SR results. Comparison curves of adaptive-scale SR performance of different methods and the SR results (corresponding to the sampling rates of 30×, 40×, and 60× objectives); Scale bars: 0.5 μm. **(c)** Deblurring: Examples of the deblur performance on the S. aureus dataset; The NRMSE values are shown on the error map images under the deblur results; Scale bar: 2 μm. **(d)** De-background: Examples of the de-background performance on mitochondria samples from DARK dataset; The NRMSE values are shown on the error map images under the de-background results; Scale bar: 2 μm. **(e)** Isotropic reconstruction: Examples of the isotropic reconstruction performance on the anisotropic mouse liver data(an axial slice); The NRMSE values are shown on the error map images under the isotropic reconstruction results; Scale bar: 5 μm. **(f)** 3D Denoising: Examples of the denoising performance of CARE, UniFMIR-Denoise, RCAN and MAGNET(with prompt) on lysosomes samples from RCAN dataset (a lateral slice and an axial slice); The NRMSE values are shown on the error map images under the denoised results; Scale bar: 1 μm. **(g)** Style transfer: Examples of the virtual fluorescence labeling performance on E.coil_membranes dataset; The NRMSE values are shown on the error map images under the style transfer results; Scale bar: 1 μm. **(h)** De-aberration: Examples of the de-aberration performance of DeAbe, UniFMIR-Deaberration, and MAGNET(with/without prompt) on light-sheet C. elegans embryos from DeAbe dataset; The NRMSE values are shown on the error map images under the de-aberration results; Scale bar: 5 μm. **(i)** 3D De-scattering: Examples of the de-scattering performance of Scatnet, UniFMIR-Descatter, RCAN and MAGNET(with prompt) on light-sheet Drosophila embryo data (a lateral slice and an axial slice); The NRMSE values are shown on the error map images under the de-scatter results; Scale bar: 20 μm. In all box plots (n = 20), the line within each box denotes the mean, and the whiskers extend to the minimum and maximum values.

In the inference phase (Fig. 1d), the trained model employs a Task Prompt Predictor (Methods) to automatically infer degradation-relevant prompts from input images, guiding the All-in-One backbone toward targeted restoration tasks. For example, for a noisy image of zebrafish embryos^28^ (Input 1), the Task Prompt Predictor automatically infers noise-related task prompts, which guide the model to perform targeted denoising, resulting in a 2.8-fold improvement in SNR relative to the raw image. For an anisotropic mouse liver volume^29^ (Input 2), the model is guided by isotropic-related prompts to reconstruct spatially consistent volumes, yielding a 2.14-fold improvement in axial resolution. Together, these results confirm that MAGNET performs adaptive, degradation-aware restoration in practice, consistently improving image quality across diverse degradation conditions.

### Multi-Task Unified Image Restoration of Cross-Dimensional Microscopy Data

In fluorescence microscopy, image degradation manifests in diverse forms, including resolution loss from limited numerical aperture (super-resolution), high noise from low excitation (denoising), and anisotropic resolution from insufficient axial sampling (isotropic reconstruction). These degradations not only obscure fine biological structures but also compromise downstream analyses such as segmentation and quantitative measurement. Addressing image degradation is therefore critical—not only to improve the quality of images but also to provide a robust foundation for quantitative biological analysis. However, most existing restoration methods are task-specific or rely on multi-head and tail foundation models with task-dependent branches, which struggle to generalize across diverse degradations without explicit fine-tuning (Fig. 2a, Supplementary Fig. 4). Moreover, multi-branch architectures often suffer from inter-task interference and limited scalability. To overcome these limitations, we propose a unified All-in-One restoration framework with a task-controllable prompt mechanism that enables adaptive recognition and targeted processing of multiple degradation types within a single network. Benefiting from this mechanism, our model effectively mitigates conflicts in multi-task learning and achieves improved generalization and structural fidelity under heterogeneous conditions. We systematically evaluated the model across a range of representative restoration tasks (super-resolution, denoising, deblurring, background suppression, isotropic reconstruction, aberration correction, de-scattering, and style transfer) to validate its broad applicability and robustness.

We first evaluated the SR capability of our method on the BioSR^16^ dataset, where SIM images serve as ground truth(GT) and WF images acquired with a 100× objective correspond to 2× down-sampled inputs. Our method accurately recovered the hollow, ring-shaped clathrin-coated pits (CCPs) with higher resolution than both DFCAN^16^ and UniFMIR, which were trained specifically on the BioSR dataset (Fig. 2b). Benefiting from the LIIF-based implicit neural representation, our model inherently supports adaptive-scale super-resolution. To further test this property, we simulated different magnification systems by downsampling the WF inputs by 3.3× (corresponding to the sampling rate of 60× objective), 5× (40× objective), and 6.6× (30× objective). Across all scales, our reconstruction consistently outperformed other methods restricted to fixed magnification training, demonstrating strong robustness and generalization for multi-scale SR tasks. Furthermore, the NRMSE curves across different downsampling scale factors confirmed that our method achieved consistently lower reconstruction errors, highlighting its superior stability and accuracy.

For deblur task, we assessed our method on a bacterial image dataset (S.aureus_MreB) from the DeepBacs^30^. Compared with CARE^29^ and UniFMIR, which were trained specifically on this dataset, our approach restored bacterial texture details with higher fidelity, achieving a 2×resolution improvement(Fig. 2c). Moreover, across synthetic blur kernels of varying orientations and magnitudes, our model maintained stable SSIM, demonstrating robustness under diverse blur conditions (Supplementary Fig. 5).

Next, we assessed background suppression performance on the DARK^31^ dataset . Our approach substantially reduced background fluorescence while preserving signal intensity, achieving a signal-to-background ratio comparable to the GT (Fig. 2d).

For isotropic reconstruction, we applied our method to anisotropic mouse liver volume^29^ (Fig. 2e). Compared with existing deep-learning approaches, our reconstructions exhibited voxel intensity distributions that more closely matched the GT, achieving near-isotropic resolution.

For denoising, tested on the publicly available RCAN^32^ 3D dataset, our approach enhanced SNR by 10×while preserving fine lysosomal structures even under extremely low photon conditions (Fig. 2f). Benefiting from the dynamic guidance of the prompt mechanism on image degradation characteristics (Supplementary Fig. 6), our model consistently outperformed other denoising approaches across a range of SNR conditions, delivering stable performance (Supplementary Fig. 6). Moreover, we further provide denoising results across different imaging modalities—including two-photon, wide-field, and confocal microscopy (Supplementary Fig. 7), demonstrating the robustness and broad applicability of our model across diverse imaging systems.

We further evaluated style transfer task, which emphasizes semantic fluorescence pattern translation. The model was trained to transfer brigh-field images of E. coil_membranes into fluorescence images from DeepBacs^30^ dataset. By incorporating content-aware prompt mechanism(Methods), our model achieved task-specific feature modulation, enabling stable style-transfer performance within multi-task training (Fig. 2g).

For aberration correction, tested on C. elegans embryo light-sheet data from DeAbe^33^, our method accurately restores cellular morphology in aberration-degraded images (Fig. 2h). We further validated its performance on simulated aberration data, confirming stable recovery across different types of aberrations. These results indicate that our approach not only corrects aberrations types seen during training but also generalizes to new imaging conditions with different aberrations (Supplementary Fig. 8).

Finally, for the de-scattering task, by using the multi-view light-sheet images of Drosophila embryos^34^ as guidance, our model effectively improves the axial resolution degraded by scattering in deep tissue (Fig. 2i). Benefiting from its 3D-aware design, the model maintained axial continuity without introducing slice-to-slice inconsistencies and achieves higher structural fidelity with improved computational efficiency compared with state-of-the-art supervised learning methods^28^ (Supplementary Table 2).

### Robust Restoration under Composite Degradations

In practical microscopy applications, images are often affected by multiple composite degradations acting simultaneously, posing significant challenges for high-quality restoration. For instance, in live-cell imaging, low excitation intensity and short exposure times are typically employed to minimize phototoxicity and photobleaching. However, these low-light conditions introduce strong noise and reduce the SNR. Additionally, the diffraction limit imposed by the limited NA of the optical system restricts spatial resolution and blurs fine structural details. And in deep-tissue imaging, light scattering and noise frequently co-occur, and their combined effects further attenuate signals and degrade image quality, making image restoration and quantitative analysis particularly challenging.

To evaluate the capability of our model in handling such complex degradations, we first simulated a series of single and composite degradation datasets by synthetically degrading clean GT images. We then trained both our All-in-One model and the current state-of-the-art multi-head and multi-tail foundation model (UniFMIR) on multiple single-degradation datasets and subsequently tested their generalization performance on unseen composite-degradation datasets (Fig. 3a). Benefiting from the unified shared-parameter design and prompt-guided task modulation, MAGNET effectively disentangled and jointly modeled multiple degradation factors, while UniFMIR—whose task-specific branches are individually optimized for single degradations— failed to capture their mutual dependencies. Even when sequential multi-stage processing (e.g., denoising followed by super-resolution) was applied, UniFMIR exhibited incomplete recovery and residual artifacts. In contrast, MAGNET achieved higher SNR and resolution more consistent with GT (Fig. 3b). Furthermore, robustness evaluations under varying noise levels (Supplementary Fig. 9) demonstrate that our model consistently outperforms other approaches in handling complex composite degradation scenarios.

**Fig. 3.**
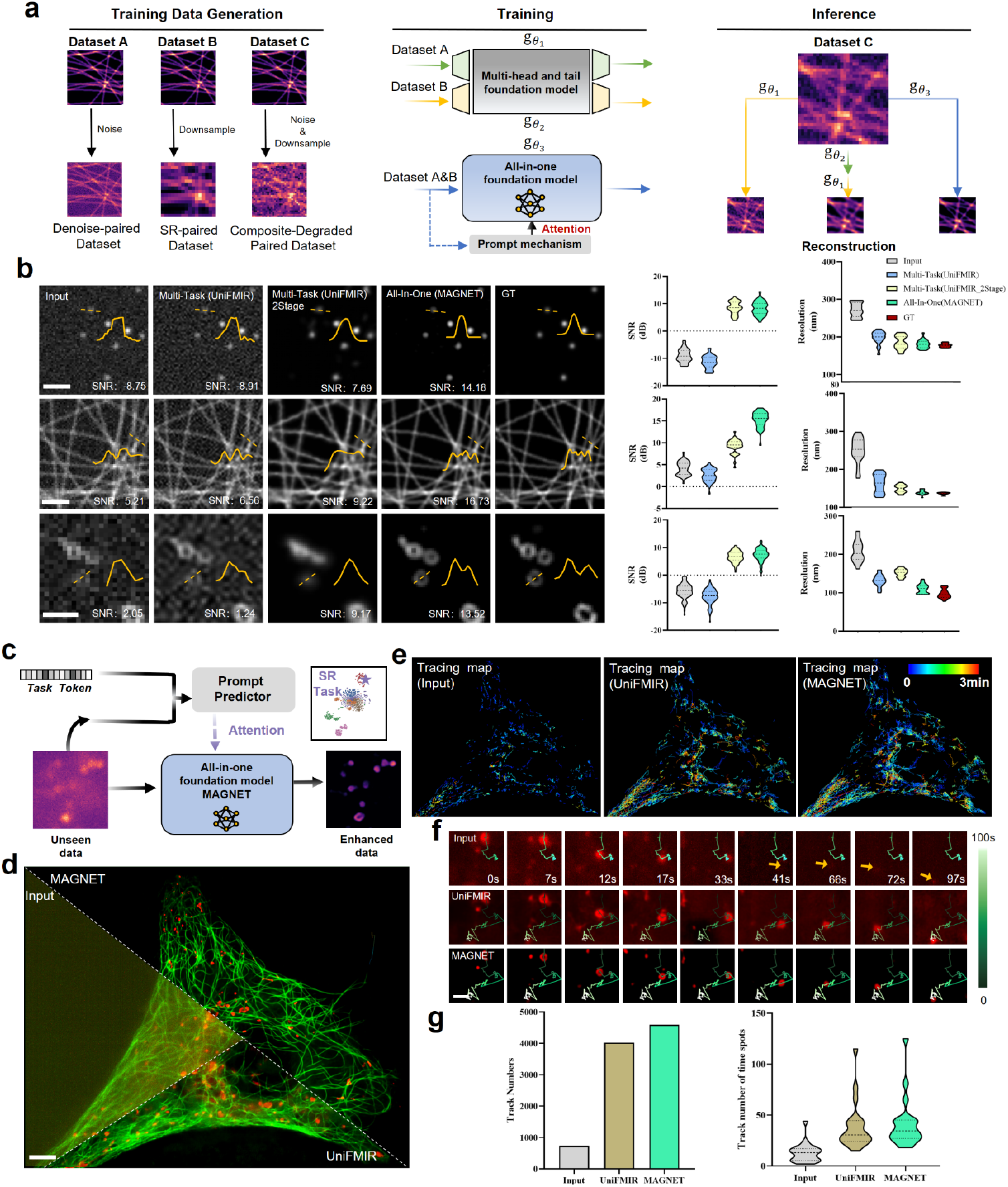
Unified restoration and downstream analysis under composite degradations. **(a)** Schematic of the generation of simulated data for different degradations, including single and composite degradations. The raw data are used as ground truth, and are then synthetically downsampled, added with noise, or subjected to both operations to produce composite degradation data; Training and inference schematic of MAGNET and the multi-head/tail foundation model. Both models are trained on single-degradation data and evaluated on composite-degradation data; For the two-stage inference approach of the multi-head/tail foundation model, the input first passes through the denoising branch and then the super-resolution branch. **(b)** Examples of the restoration performance of Multi-Task(UniFMIR), Multi-Task(UniFMIR_2Stage), and All-In-One(MAGNET) on simulated data (spots, filaments, and rings) under unseen composite degradations; SNR*↑* (higher is better)/Resolution*↓* comparisons of different methods on different datasets; Scale bars: 1 μm. **(c)** Schematic of the Task Prompt Predictor, which generates task-related prompt embeddings directly from the input image and task token. **(d)** Comparison of image restoration performance on maximum-intensity projections (MIPs) of dual-channel live-cell imaging data (microtubules tagged with 3×mEmerald-ensconsin and lysosomes tagged with Rab7-EGFP or Rab7-mCherry) acquired using a custom line-scanning confocal microscope; Scale bar: 10 μm. **(e)** Comparison of lysosome tracking results based on raw input, UniFMIR-restored, and MAGNET–restored images; The colored curves in the figure indicate the time-coded trajectories. **(f)** Example of single lysosome tracking results; Yellow arrows indicate trajectories that are not captured in the low-SNR raw inputs; The colored curves indicate the time-coded trajectories; Scale bar: 1 μm. **(g)** Statistical comparison of the number of detected trajectories and the track durations. In box plots of g (n = 20), the line within each box denotes the mean, and the whiskers extend to the minimum and maximum values.

We next evaluated whether our model could generalize to real-world composite degradations and improve downstream live-cell analysis tasks, such as lysosome tracking. To address the challenge of missing paired prompt images during inference, we designed a Task Prompt Predictor (Fig. 3c, Methods, and Supplementary Fig.10), which enhances the model’s adaptability under unknown degradations by generating task-related prompt embedding directly from the input image and corresponding task token. Applying both our pre-trained model and UniFMIR to dual-channel live-cell datasets (microtubules tagged with 3×mEmerald-ensconsin and lysosomes tagged with Rab7-EGFP or Rab7-mCherry) acquired using a custom line-scanning confocal microscope (Methods), our model demonstrated robust performance even on previously unseen, extremely noisy real-time imaging data (Fig. 3d). To assess the benefits for cell tracking, we applied a simple tracking pipeline to the raw, UniFMIR-restored, and our restored image stacks (Fig. 3e). Our method not only recovered the hollow ring-shaped lysosomes more accurately but also suppressed noise more effectively, thereby improving tracking precision (Fig. 3f). Furthermore, the increased number of detected trajectories and longer average tracking durations indicate that our restored data supports high-speed and long-term imaging with enhanced reliability, facilitating improved cell lineage tracking and dynamic biological analysis (Fig. 3g).

### Progressive Deployment enables Flexible Generalization across Diverse Imaging Conditions

In practical microscopy, the sources of image degradation are highly system-dependent and non-ideal, manifesting as complex statistical variations introduced by differences in imaging system configurations, sample types, noise levels, and experimental conditions. Although existing deep learning–based image restoration methods often achieve impressive performance on specific datasets, they typically lack sufficient adaptability and generalization robustness when confronted with unseen imaging systems, unknown degradation types, or out-of-distribution samples. This limitation hinders their applicability to the diverse experimental requirements of real-world scenarios. Moreover, acquiring high-quality paired training data is costly—particularly for biological samples under low SNR conditions or during dynamic processes—posing a practical bottleneck for traditional supervised learning paradigms that rely on large-scale annotated datasets.

An ideal image restoration framework should therefore be capable of direct inference(out-of-the-box) without additional task-specific training, efficient adaptation under limited annotated data (few-shot fine-tuning), and self-supervised test-time optimization when annotations are unavailable (zero-shot test-time optimization). Such capabilities are crucial for dealing with the inherently variable and uncontrollable degradation patterns encountered in microscopy imaging.

To evaluate cross-modality generalization on seen samples (Out-of-the-box direct inference), we directly applied our pretrained model—without any retraining or fine-tuning—to microtubule and endoplasmic reticulum (ER) datasets acquired by our custom-built line-scanning confocal and light-sheet fluorescence microscopes (Fig. 4a). Despite the challenging presence of complex composite degradations in the test data—including low-resolution images with pronounced noise and images corrupted by combined blur and noise—our proposed All-in-One model consistently demonstrated superior structural restoration and noise suppression compared to other deep learning– based SR methods across multiple unseen test datasets, highlighting its strong generalizability.

**Fig. 4.**
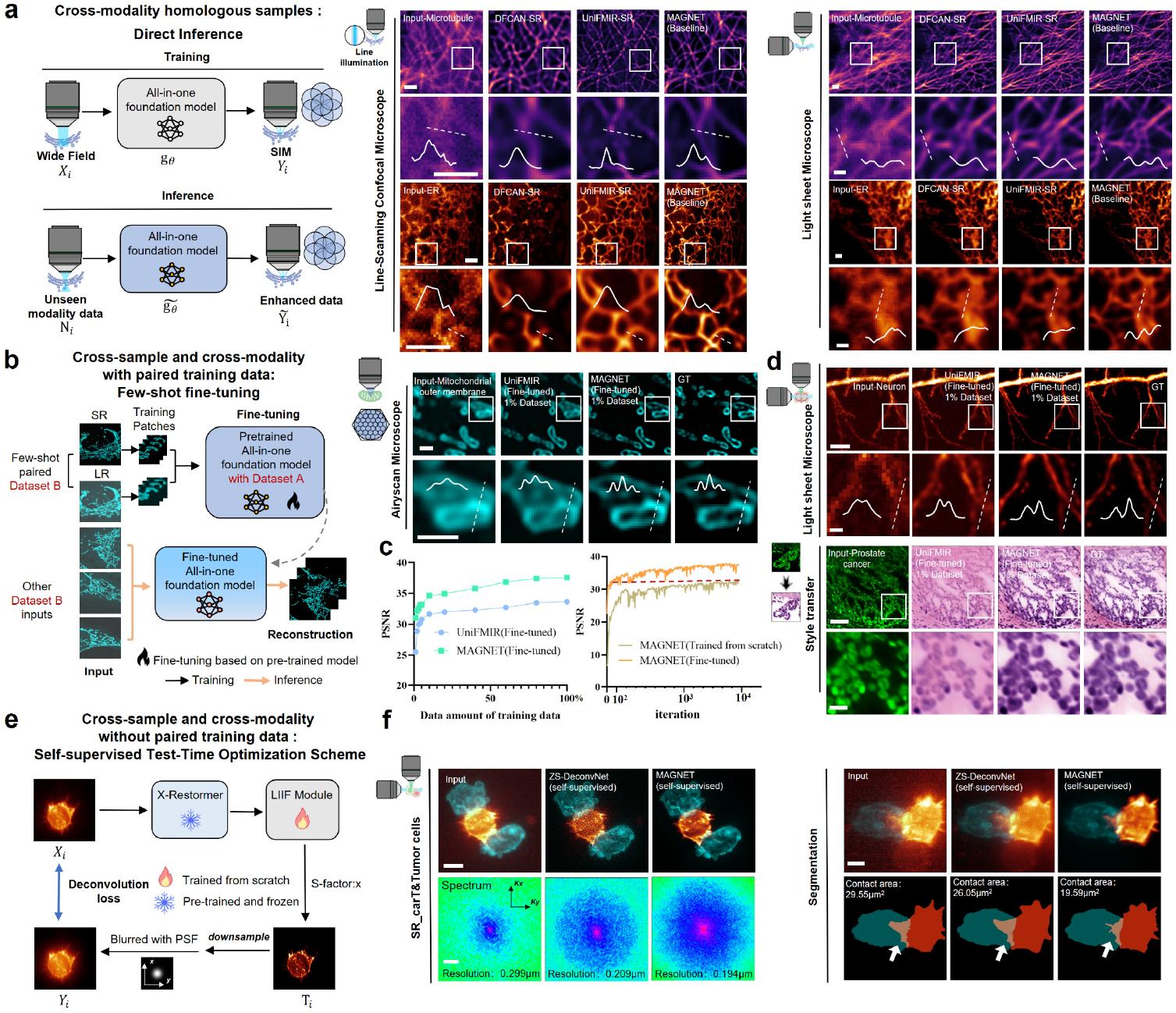
Generalization capability enabled by progressive deployment of MAGNET. **(a)** Cross-modality direct inference(Out-of-the-box); Examples of the restoration performance of DFCAN-SR, UniFMIR-SR, and MAGNET(Baseline) on homologous datasets of microtubules and endoplasmic reticulum (ER) acquired using custom-built line-scanning confocal and light-sheet microscopes without retraining or fine-tuning (a lateral slice). Scale bars: 1 μm. **(b)** Schematic of few-shot fine-tuning with paired training data; The pretrained MAGNET (trained on large-scale datasets) is further fine-tuned using a small number of paired samples; Examples of the restoration performance of UniFMIR(Fine-tuned) and MAGNET(Fine-tuned) on mitochondria outer membrane images(a lateral slice) acquired by the Airyscan super-resolution microscope(both sample type and imaging system are unseen); Scale bars: 1 μm. **(c)** Comparison of model performance under varying sample proportions and training strategies. **(d)** Few-shot fine-tuning results of UniFMIR(Fine-tuned) and MAGNET(Fine-tuned) on other cross-sample and cross-modality datasets—including light-sheet mouse brain nerve images and fluorescence-to-H&E prostate cancer tissue images(a lateral slice); Scale bars from top to bottom: 10 μm, 2 μm, 100 μm, 20 μm. **(e)** Schematic of the MAGNET self-supervised test-time optimization (TTO) scheme; In the absence of ground-truth data, the system optical PSF and low-quality images are used for self-supervised optimization, where the X-Restormer backbone is kept frozen and the LIIF-based image reconstruction module is fine-tuned. **(f)** Examples of GT-free restoration results of ZS-DeconvNet (self-supervised) and MAGNET (self-supervised) on maximum-intensity-projection (MIP) images from light-sheet microscopy data of CAR-T and tumor cell interactions. Fourier spectra are shown to quantify spatial resolution. Cell segmentation results using Cellpose^39^ are shown, with white arrows indicating interaction regions of CAR-T and tumor cells. Scale bars: 5 μm.

In practical microscopy applications, although paired training data can be obtained in some scenarios, the acquisition of high-quality paired samples is often challenging to obtain—particularly for multimodal imaging (e.g., fluorescence-to-H&E), SR modalities (e.g., Airyscan super-resolution microscope^35^), or deep tissue imaging. These scenarios frequently encounter technical barriers such as limited fields of view, photobleaching, and complex imaging setups, which impede the construction of large-scale training datasets. Consequently, large-scale supervised training becomes an unrealistic goal. Against this backdrop, leveraging existing domain knowledge as cross-domain priors and achieving efficient adaptation with limited samples emerges as a critical practical strategy. To assess cross-sample and cross-modality adaptability with paired training data (Few-shot fine-tuning), we evaluated our model on mitochondria outer membrane images acquired by Airyscan super-resolution microscope^35^ (Fig. 4b). In cases where both sample types and imaging systems are unseen, direct inference performance typically degrades. However, with lightweight fine-tuning on a small number of paired samples, our model’s restoration accuracy improved significantly. To further assess the model’s stability under few-shot conditions, we progressively reduced the training set and plotted the relationship between sample proportion and restoration performance. Results demonstrate that even when training samples are reduced to 1%, the model maintains stable performance without significant degradation, underscoring its superior robustness and adaptability in low-data regimes. To further quantify adaptation efficiency, we compared the performance curves of our model fine-tuned from pretrained weights versus trained from scratch. Results show that, benefiting from large-scale pretraining, our model achieved high restoration accuracy within a fraction of the training time required for full retraining—demonstrating substantial efficiency gains (Fig. 4c). Beyond mitochondria restoration, MAGNET consistently demonstrates strong cross-sample and cross-modality adaptability across additional tasks (Fig. 4d). For instance, on light-sheet mouse brain neuron dataset (Methods), few-shot fine-tuning enables MAGNET to recover markedly clear neuronal morphology, revealing fine dendritic structures. Similarly, in fluorescence-to-H&E style transfer, our model produces morphologically consistent tissue structures and more accurate prostate cancer lesion depiction, achieving pathology-grade contrast that closely matches the GT^36^ (Supplementary Fig.11).

In certain biological application scenarios, such as long-term live-cell dynamic imaging or neural signal transmission recording, acquiring GT images or paired training samples is difficult. To achieve high-quality image restoration without paired supervision, we propose a self-supervised test-time optimization (TTO) scheme (Methods) built upon our multi-task pretrained model, and validate it on light-sheet microscopy datasets^37^ capturing CAR-T and tumor cell interactions. Without any GT images, we exploit only the system’s optical PSF prior and the original low-quality images for self-supervised training. Specifically (Fig. 4e), we freeze the X-Restormer backbone and fine-tune only the LIIF module parameters. The training adopts a single-stage optimization strategy, integrating the system’s measured or simulated PSF to enforce physical consistency and enhance structural fidelity. Compared with the state-of-the-art self-supervised learning method ZS-DeconvNet^38^, which relies on optimization from limited test data, our model pretrained on large-scale, diverse tasks and datasets. Therefore, our model yields reconstructions with reduced residual noise, and improved resolution even without GT supervision (Fig. 4f). Enhanced resolution facilitates downstream quantitative analyses—enabling more accurate segmentation of CAR-T and tumor cells and precise measurement of their contact interfaces—thus demonstrating strong potential for achieving high-fidelity image restoration under GT-free constraints (Fig. 4f).

## Discussion

In this study, we introduced MAGNET, an All-in-One foundation model that unifies 2D and 3D microscopic data and accommodates diverse degradation types within a single framework. By combining a triple-plane projection module, a prompt-guided X-Restormer backbone and an LIIF-based continuous scale reconstruction module, MAGNET supports multi-task, cross-modal and resolution-adaptive restoration in a single framework. To support large-scale multi-task training, we collected a substantial microscopic image restoration dataset, including more than 10 public datasets together with multiple in-house datasets. The resulting corpus contains tens of thousands of image pairs, spans 5+ major imaging modalities (confocal, wide-field, SIM, light-sheet, and two-photon), and reaches a total size of over 500 GB, forming the most extensive and diverse training resource to date for microscopic image restoration. Evaluated across 8 representative tasks and more than 5 major microscopy modalities, the model consistently achieves state-of-the-art performance across diverse imaging conditions. Benefiting from its unified representation and adaptive prompt guidance, MAGNET not only excels in single degradation tasks—such as super-resolution (achieving robust performance across variable scales by elevating the effective sampling rates associated with 30×∼100× objectives to levels approaching the *∼*200× sampling regime of SIM.)— but also demonstrates superior robustness under composite degradations. Moreover, its progressive deployment paradigm, encompassing out-of-the-box direct inference, few-shot fine-tuning, and self-supervised test-time optimization, ensures high flexibility and adaptability across diverse experimental settings and data availabilities.

Beyond methodological contributions, MAGNET has practical implications for biological imaging. Higher-resolution and denoised reconstructions can facilitate more precise analysis of organelle morphology and dynamics, such as mitochondrial fission-fusion and lysosome transport; robust super-resolution in thick tissue improves delineation of fine neuronal processes; and fluorescence-to-H&E style transfer may help bridge molecular contrast with histopathological interpretation in high-throughput workflows. We anticipate that such applications will make All-in-One restoration models increasingly relevant to routine quantitative imaging.

Despite these advances, challenges remain. Highly complex or coupled degradations still pose ill-posed inverse problems, where faithful recovery remains limited by data diversity and physical priors. Additionally, with the continuous emergence of new imaging modalities and acquisition technologies, the adaptability of current architectures to unseen modalities remains uncertain. Future research will investigate the incorporation of generative models^40^, exploiting their ability to capture the statistical distribution of high-fidelity images, to improve image restoration performance under challenging, composite degradation conditions. Another promising direction is the integration of temporal dynamics to unify spatial and temporal restoration, enabling the 4D reconstruction of high-speed biological processes. Finally, bridging low-level image restoration and high-level semantic reasoning—by coupling large-scale vision-language models or biological priors—may open new avenues toward interpretable, intelligent microscopy.

Overall, MAGNET establishes a powerful foundation model for microscopy by unifying heterogeneous degradation types, imaging modalities, and data dimensionalities within one architecture. This unified design enhances the universality, robustness, and flexibility of microscopic image restoration. Beyond improving image visual quality, MAGNET serves as a general computational tool to accelerate biological discovery—providing cleaner, more reliable data for downstream analyses across multiple applications. We anticipate that this unified paradigm will inspire future research on foundation models, driving the deeper integration of AI with biological imaging.

## Methods

### Prompt-based task disentanglement

A central challenge in All-in-One microscopic image restoration lies in the pronounced heterogeneity across tasks, which makes unified models prone to catastrophic forgetting and inter-task interference, resulting in degraded performance. In practice, many restoration networks that excel in one task fail to generalize to others, underscoring the difficulty of achieving true task generality. To address this, we adopt X-Restormer^41^ as a task-general backbone and introduce a task prompt mechanism with an accompanying Task Prompt Predictor, enabling effective task disentanglement and adaptive restoration within a unified framework.

### Task prompts mechanism

Prompt learning was originally proposed in the field of natural language processing (NLP)^42^, where manually crafted context prompts were supplied to pretrained models to perform downstream tasks without fine-tuning the backbone parameters. Subsequently, the paradigm was extended by replacing handcrafted prompts with task-specific learnable vectors^43, 44^, thereby greatly improving adaptability and generalizability. More recently, prompt learning has been introduced into computer vision^45, 46^, where it has proven increasingly effective for unifying diverse vision tasks.

In our design, we adopt this concept for unified microscopy image restoration tasks. Specifically, these tasks are embedded into an “task semantic space”, and paired prompt latent vectors (prompt embeddings) are introduced to guide the backbone network in adaptively adjusting its computational pathway (Supplementary Fig. 3b).

Given an input image I_*in*_, high-dimensional semantic features Z_*in*_ are first extracted through the encoder of the backbone. At the same time, a pair of prompt images [P_*input*_, P_*target*_] is fed into a dedicated Task Prompt Encoder, which outputs prompt semantic features 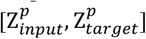 aligned in spatial dimension with Z_*in*_. To enhance semantic interaction between prompt features and image representation, we further design a Prompt Cross-Attention Block (PCAB) and embed it at multiple key stages of the backbone. In PCAB, the query(Q), key (K) and value (V) are first generated by 1 × 1 convolutions from the input representation Z_*in*_, prompt input embedding 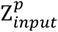 and prompt target embedding 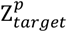. Then, the standard attention is computed to obtain the output representation Z_*out*_, which serves as the deep semantic representation for downstream image restoration. The cross-attention operation is formulated as:

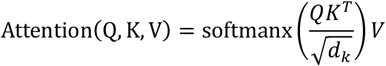

### Task prompt predictor

In practical microscopy, obtaining paired prompt images is often infeasible, particularly during inference and in zero-shot scenarios. To address this limitation, we designed a Task Prompt Predictor that directly predicts a latent prompt representation from the input image itself, thereby guiding task-specific modeling and image restoration without requiring explicit paired prompts (Supplementary Fig.10). Specifically, the module takes as input the high-dimensional feature representation Z_*in*_ together with its task encoding E_*task*_. These inputs are jointly mapped through a lightweight network into a task-relevant latent prompt representation Z_*pred*_, which serves as a surrogate for the paired prompt features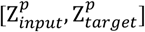 (e.g., a one-hot vector or a learnable task prompt embedding) in the Prompt Cross-Attention mechanism. The Predictor is implemented using a standard CNN-based feature extractor, and trained on a synthetic dataset. This dataset incorporates representative biological structures—including punctate structures (e.g., nuclei or vesicles), filamentous structures (e.g., cytoskeletal fibers), and ring-like structures (e.g., circular organelle boundaries)—subjected to diverse degradations such as noise, blur, and low resolution. Each degraded sample is paired with a ground-truth latent prompt embedding, and the Predictor is trained by minimizing the distance (e.g., L2 loss or cosine similarity) between the predicted prompt embedding Z_*pred*_ and the ground-truth prompt embedding Z_*out*_. Through this strategy, the module learns to generate task-relevant latent prompts under real experimental conditions where explicit paired prompt images are unavailable.

### Projection-based dimensionality adaption

Microscopy data span heterogeneous spatial dimensionalities, ranging from single 2D images to volumetric 3D stacks, posing a fundamental challenge for unified modeling within a unified restoration framework. Directly mixing 2D and 3D representations can lead to semantic conflicts and inefficient feature sharing. To address this, we introduce a projection-based dimensionality adaptation strategy that converts heterogeneous inputs into dimension-compatible representations while preserving essential structural and textural information.

### Triple-plane projection

To address the challenge of integrating heterogeneous microscopy data encompassing both 2D images and 3D volumes, we introduced a triple-plane projection module into the model (Supplementary Fig. 3a). This module enables the network to process both 2D and 3D inputs and to derive dimension-compatible feature representation. For 2D images, a single mean projection is computed to generate the initial feature representation. For 3D volumes, projections are computed along the x, y, and z axes, and three statistical descriptors—mean, maximum, and variance—are computed along the channel dimension to generate complementary 2D feature maps (Supplementary Fig.12). This design is inspired by prior study from medical imaging^47^ and remote sensing^48^, showing that statistical projections can effectively preserve textural and structural information in multi-channel data. The projected multi-view 2D features are then fed into the X-Restormer backbone for deep feature enhancement. Finally, to recover the volumetric structural information intrinsic to the original 3D volumes, we introduce a 3D reconstruction module (Supplementary Fig. 3c) that transforms the enhanced 2D representations back into enhanced 3D feature volumes, thereby achieving dimensional-compatible feature enhancement and information restoration.

### Dimensionality-adaptable feature fusion

To enhance the network’s ability in handling inputs of different spatial dimensionalities and to mitigate potential semantic conflicts during cross-dimensional modeling, we introduce a dimension prompt mechanism into the architecture (Supplementary Fig.13). Unlike the task prompt mechanism described earlier, which primarily encodes image degradation semantics, the dimension prompt mechanism explicitly conveys dimensionality information by injecting structured dimension-embeddings into the feature stream. In this way, the network gains an enhanced ability to perceive and adapt to dimensional differences.

Specifically, we insert dimension embeddings at two critical positions: (1) immediately after the triple-plane projection module and before the X-Restormer backbone, where they explicitly distinguish the geometric structures corresponding to 2D versus 3D inputs and guide the backbone to adjust its attention modeling strategy accordingly; and (2) after the X-Restormer backbone and before the LIIF image reconstruction module, where they provide dimensionality cues during the continuous scale reconstruction process, thereby modulating the feature fusion pathway during querying.

In implementation, we define a dimensionality indicator variable task_*id*_ ∈ {0, 1}, corresponding to 2D and 3D inputs, respectively. A set of learnable dimension embeddings and modulation parameters (task_*y*_ and task_*p*_) are then fused with the network features, enabling conditional normalization and feature guidance under different spatial dimensionalities. This design provides the network with explicit awareness of spatial dimensional properties, effectively alleviating representational conflicts between 2D and 3D inputs and establishing a structural foundation for unified cross-dimensional modeling.

### LIIF-based continuous scale restoration

To enable continuous and resolution-adaptive image restoration across diverse imaging conditions, we introduce a dimension-adaptable implicit representation module inspired by LIIF^49^. By extending implicit neural representations to accommodate different spatial dimensionalities and microscopy-specific sampling characteristics, this module supports flexible, resolution-continuous reconstruction within a unified framework. In the context of implicit image reconstruction, existing LIIF-based approaches are typically designed for 2D images only, lacking the capacity to model continuous structures in 3D space. This restricts their applicability to volumetric imaging tasks. To address this, we extend the LIIF framework by designing a dimension-adaptable 2D/3D implicit image reconstruction module, which enables unified modeling of both 2D pixels and 3D voxels within a shared network architecture(Supplementary Fig. 3d).The module incorporates a dimension-adaptable coordinate querying mechanism that flexibly adapts feature sampling and reconstruction to the spatial dimensionality of the input. In the 2D mode, we follow the original LIIF querying strategy: for each continuous coordinate, its representation is obtained through weighted interpolation of the surrounding 2×2 discrete feature points. In the 3D mode, this process is naturally extended to voxel space, where the representation at a continuous 3D location is computed by weighted interpolation of features from the neighboring 2×2×2 voxel points. This extension ensures accurate and coherent volumetric reconstruction, thereby enabling high-fidelity restoration across both 2D and 3D microscopy data.

### Progressive deployment strategy

In real microscopy applications, restoration models must operate under heterogeneous data forms and degradation conditions. To accommodate this variability, we design a progressive deployment strategy for MAGNET encompassing out-of-the-box direct inference, few-shot fine-tuning with limited paired data, and self-supervised test-time optimization(TTO) in the absence of ground truth.

To address scenarios where ground-truth supervision is unavailable, we implemented a self-supervised TTO strategy on top of the pretrained model. During TTO, only the lightweight MLP layers in the LIIF are updated, while all other parameters remain frozen, thereby minimizing overfitting risk. Following the principle of ZS-DeconvNet^38^, we designed a self-reconstruction training using the system point spread function(PSF).

Unlike existing self-supervised approaches such as Noise2Noise^50^, which rely on dual U-Net architectures, our design integrates a LIIF-based implicit image representation module, which decouples the super-resolution scale from the network architecture. This allows a single model to flexibly perform arbitrary-scale super-resolution reconstruction. The reconstructed output at an arbitrary continuous spatial coordinate is expressed as:

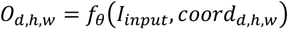

where *I*_*input*_ is the degraded image, *d, h, w* denote spatial indices, and *coord* encodes the continuous coordinate query. Then the measured or simulated point spread function (PSF) is incorporated to further refine structural clarity:

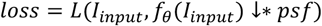

where *f*_*0*_(·) denotes the LIIF reconstruction module with trainable parameters *θ, ↓* indicates downsampling, and *\** represents convolution.

### Data acquisition

For real-world composite degradations experiment, the dual-channel live-cell datasets (microtubules tagged with 3×mEmerald-ensconsin plasmid and lysosomes tagged with Rab7-EGFP plasmid or Rab7-mCherry plasmid) were acquired by a custom line-scanning confocal microscope. An oil immersion objective (UPLSAPO100X, 100×/1.4NA, Olympus) was used to collect the epifluorescence signals from samples with scientific camera sensor (Flash 4.0 V2, Hamamatsu). The excitation source of the system is a laser with wavelengths of 488nm and 561nm.

For few-shot fine-tuning experiment, the mouse brain neuron dataset(transgenic line: Thy1-GFP) was acquired by a custom light-sheet microscope. An oil immersion objective (UPLXAPO40XO, 40×/1.4NA, Olympus) was used to collect the epifluorescence signals from samples with scientific camera sensor (Flash 4.0 V2, Hamamatsu). The excitation source of the system is a laser with wavelength of 488nm.

## Acknowledgements

We are grateful to Yao Zhou, Jie Wang, Dr. Fang Zhao and Dr. Minglu Sun for providing us the fluorescence and pathology dataset. We are grateful to Dr. Yunzhe Li for helpful suggestions and for editing the manuscript. This work was supported by the fundings from National Natural Science Foundation of China (T2225014, 22534002, 62505099), National Key Research and Development Program of China (2022YFC3401100, 2023ZD0519900, 2025YFF1500200), China Postdoctoral Science Foundation (2025M780778), Key Research and Development Project of Hubei Province (2024BCB112), and The Interdisciplinary Research Program of HUST(YCJJ20252104).

## Author contributions

P.F., Y.M. and B.L. conceived the idea. P.F., and B.L. oversaw the project. Y.M., C.Y., and T.Z. developed the programs. Y.M., T.Z., and Y.Z. processed the images. Y.M., L.Z., B.L. and P.F. analyzed the data and wrote the paper.

## Notes

### Competing Interest Statement

The authors have declared no competing interest.

